# Electroceutical disinfection strategies impair the motility of pathogenic *Pseudomonas aeruginosa* and *Escherichia coli*

**DOI:** 10.1101/088120

**Authors:** Kristina Doxsee, Ryan Berthelot, Suresh Neethirajan

## Abstract

Electrotaxis or galvanotaxis refers to the migration pattern of cells induced in response to electrical potential. Although it has been extensively studied in mammalian cells, electrotaxis has not been explored in detail in bacterial cells; information regarding the impact of current on pathogenic bacteria is severely lacking. Therefore, we designed a series of single and multi-cue experiments to assess the impact of varying currents on bacterial motility dynamics in pathogenic multi-drug resistant (MDR) strains of *Pseudomonas aeruginosa* and *Escherichia coli* using a microfluidic platform. Motility plays key roles in bacterial migration and the colonization of surfaces during the formation of biofilms, which are inherently recalcitrant to removal and resistant to traditional disinfection strategies (e.g. antibiotics). Use of the microfluidic platform allows for exposure to current, which can be supplied at a range that is biocidal to bacteria, yet physiologically safe in humans (single cue). This system also allows for multi-cue experiments where acetic acid, a relatively safe compound with anti-fouling/antimicrobial properties, can be combined with current to enhance disinfection. These strategies may offer substantial therapeutic benefits, specifically for the treatment of biofilm infections, such as those found in the wound environment. Furthermore, microfluidic systems have been successfully used to model the unique microfluidic dynamics present in the wound environment, suggesting that these investigations could be extended to more complex biological systems. Our results showed that the application of current in combination with acetic acid has profound inhibitory effects on MDR strains of *P. aeruginosa* and *E. coli*, even with brief applications. Specifically, *E. coli* motility dynamics and cell survival were significantly impaired starting at a concentration of 125 μA DC and 0.31% acetic acid, while *P. aeruginosa* was impaired at 70 μA and 0.31% acetic acid. As these strains are relevant wound pathogens, it is likely that this strategy would be effective against similar strains *in vivo* and could represent a new approach to hasten wound healing.

## Introduction

Antibiotic resistance, although a long-standing issue, continues to be a growing problem. The number and prevalence of multi-drug (MDR) resistant bacterial strains is steadily increasing, and more than 2 million people are infected each year in the United States by resistant strains, causing some 23,000 deaths^1^. An important contributor to antibiotic resistant infection is biofilm infection; by definition biofilms are inherently resistant to antimicrobials, warranting up to 1000 times the dose required to clear non-biofilm infections^2^. Therefore, alternative strategies to disinfection (e.g. electrocidal, electroceutical, and mechanical approaches) have gained interest and represent potentially powerful weapons against recalcitrant biofilm infection. Although the impact of current on mammalian cells is fairly well understood, information regarding the effect on pathogenic MDR bacterial strains is lacking.

A number of studies have investigated bacterial responses to electrical fields and applied current. For example, the motility of *E. coli* cells has been evaluated in capillaries as a facet of soil remediation initiatives^3^, and *Tetrahymena pyriformis* electrotactic behaviors have been investigated with the goal of developing a microrobot^4^. The efficacy of electrical stimulation as a means of bacteriostatic control in the areas of wound healing^5^^6^^7^^8^^9^and biofilm remediation^10^^11^^12^^13^ has also been investigated in recent years. However, data on the cellular motility and electrotactic and galvanotactic behaviors MDR bacterial species (in response to the external application of current) is rather limited. Electrical stimulation, particularly in the 50-500 μA direct current (DC) range, has been shown to reduce the number of viable cells, as determined by colony forming units (CFUs). However, the underlying mechanisms by which electrical currents exert these effects in bacterial cells are poorly understood. Not only can current affect viability, but it can also impact the motility of bacterial organisms. However, the effect of current on the motility of pathogenic bacteria has not been extensively explored.

Cell migration/motility plays important roles in wound healing. Diverse cellular factors, such as chemical and electrical cues, guide the directional migration of several organisms^14^. For example, leukocytes (e.g. lymphocytes and neutrophils) can detect and follow gradients of tissue-derived chemoattractants^15^ and electric fields generated endogenously at wound sites to foster healing and antimicrobial defense^16^.Therefore, the strategic application of current could augment natural wound healing responses in mammalian cells, while targeting pathogenic bacteria. MDR strains of *Pseudomonas aeruginosa* and *Escherichia coli* are of particular interest in the wound environment, as they are common wound pathogens that possess motility systems critical to biofilm formation; directed motility towards a surface and eventual loss of the flagella are essential steps in biofilm formation in these organisms^17^. *Pseudomonas aeruginosa*, a major cause of nosocomial infections, frequently infects open wounds and can cause sepsis and necrosis^18^. This organism infects approximately one-third of all burn wounds^19^. Additionally, *Pseudomonas aeruginosa* has negative industrial and environmental influences, causing dairy spoilage^20^ and issues in water treatment systems^21^. Therefore, applying electric potential could be a complementary approach in addition to using chemical-based drugs to overcome the issue of multi-drug resistance. Novel electroceutical technologies incorporating low voltage/current within a wound dressing have shown some promise and could be implemented as a wearable electrical-based treatment system^22^. Therefore, understanding electrotaxis behaviors in MDR strains of bacteria will provide researchers with relevant information concerning the ranges of electric potential that could be applied to chronic or acute wound infections, depending on the infecting strains and their relative sensitivities.

Such electroceutical strategies combine chemical disinfection with the application of current. Acetic Acid (AA) is one promising chemical candidate that may be effective in such strategies, as it is an effective, safe, and economical antimicrobial agent capable of inhibiting pathogenic and MDR strains of bacteria, even when these strains grow as a biofilm^19^,^23^. Although studies strongly support the use of AA in combination with electrical stimulation (ES) as a disinfection strategy, additional measurement of the chemotactic and electrotactic behaviors of MDR strains is needed to engineer and fabricate an effective wound healing device. Therefore, we chose to compare the effects of AA or ES alone (single cue) or a combination of AA and ES (multi-cue) on the chemotaxis of pathogenic species relevant to wound infection.

Our results show that combinations of AA and ES are highly effective in reducing the motility of MDR strains of *P. aeruginosa* and *E. coli*, which is likely to impede infection and biofilm formation. Our results help to deepen our understanding of the effects of electroceutical approaches on pathogenic bacteria, suggesting that motility is one heavily impacted factor. This information may aid in the design of highly effective wound healing devices/strategies.

## Materials and Methods

### Experimental Parameter Design

Care was taken in the selection of various experimental parameters, which will be discussed herein. Studies support that both *E. coli* and *P. aeruginosa* show little to no response to alternating current (AC) stimulation, but a bacteriostatic effect is observed when exposed to anodal and cathodal direct current (DC)^7^,^24^. Delivery of 200 μA for 4 h/day over 4 days reduced *P. aeruginosa* biofilms on Teflon and titanium discs^12^, and exposure to 100 μA resulted in observable biofilm reductions after 4 days^18^. Further, *in vivo* studies have shown that skin ulcers colonized with *P. aeruginosa* and treated with μA cathodal DC resulted pathogen-free ulcers within days of treatment^16^. Additional studies revealed that delivery of current through carbon-filled electrodes to microorganisms in intact human skin at 75 and 100 μA resulted in bactericidal effects at 4 and 24 hours, beneath the positive electrode^24^. We selected *P. aeruginosa* strain BK-76, isolated from canine ear skin infections, and *E. coli* strain ATCC 8099, because these are relevant wound pathogens that exhibit antimicrobial resistance.

To minimize the risk of pH or temperature fluctuations, while selecting an effective current range, we elected to administer 75, 125, or175 μA DC to cells, as that doses of 75-200 μA have bactericidal effects. Based on a study performed at the Mayo Clinic, the deployment of low dose electric current in the urinary tract was determined to be safe; a study of electrified catheters in sheep resulted in no chemical or physical changes/trauma to tissues or urine within the urinary tract when administering 400 μA of current. A similar effect was observed in a human population using an electrified urinary tract catheter trial^9^. However, it is unknown as to how inflamed and possibly necrotized skin wounds would respond to currents as great as 400 μA DC, which is why we selected a lower 175 μA dose as the upper limit. The bacterial suspension consisted of either *P. aeruginosa* or *E. coli* suspended in filtered deionized water. This was done to reduce the complexity of the electrochemical products produced at the anode/cathode. This allowed for a more clear observation of the bacterial response to current in terms of their chemotactic behavior. When the bacteria were observed microscopically in the prepared solution, they exhibited a high level of free motility.

Previous studies have shown that a minimum inhibitory concentration (MIC) of 0.31% Acetic Acid to be an effective treatment against a multitude of pathogens tested, including MDR strains of *E. coli* and *P. aeruginosa*^19^. This same concentration of AA was also found to be effective in inhibiting biofilm formation^19^. Therefore, we chose to us 0.31% as the chemical component of our electroceutical approach (multi-cue experiments).

Electrotaxis and chemotaxis have been largely been studied separately due to the complications in designing simultaneous chemical mixing and electrical field gradients in a single device^14^. The development of specialized microfluidic devices have allowed researchers to test impact of electric fields as well as controlled chemical gradients, although still challenging for practical application in a wound dressing^14^. Therefore, we elected to use a single uniform concentration of AA in combination with the application of various currents.

### Cell Culture

Bacterial suspensions were prepared by centrifuging cultures grown in 5 mL of Tryptic Soy Broth medium in the shaker at 200 rpm for 5 h at 37°C. The media was extracted and centrifuged at 3750 rpm for 5 min to concentrate the cells. Upon pouring off the supernatant and redistributing the cells in filtered deionized water, the process was repeated 2 times. The cells were subsequently diluted with filtered deionized water and injected into the microfluidic device for viewing. The above procedure was also followed for the culturing of the cells for the electroceutical experiments, except 6.9 μL of 45% Acetic Acid was added to 993.1 μL of the bacterial solution to achieve the 0.31% AA concentration prior to injection into the microfluidic channel.

### Metrics Acquisition

Copper electrodes (diameter, 4 mm) were inserted into ports A & B and into the bacterial suspension of the glass bottom fused silica microfluidic system (Figure 1). The desired ampere ranges were achieved by use of a current amplifier (Figure 1) and at the following progression and exposure times: I = 0 mA (10 min), 3 min of rest, 0.07 mA (10 min), 3 min of rest, 0.125 mA (10 min), 3 min of rest and 0.175 mA (10 min). A Nikon Ti-U Eclipse Microscope (Nikon Instruments Inc., Melville, NY) was used to image the cells at the following settings: Phase 2 contrast with the use of D & GIF filters, 40x Objective with collar ring set to 1.3 mm, 1.5X magnification, 1.0 Gain, recorded at 90 fps by use of the Nikon NIS-Elements software for real time imaging. Regions along the entire microfluidic channel were imaged and recorded for 40 seconds. The use of a flow-free microfluidic device composed of a fused silica chip, minimizes flow-induced shear stress on cell migration and movement in a static gradient environment^26^^27^^28^^29^ and allows quantitative evaluations of cell migration in spatiotemporally complex chemoattractant fields that mimic *in vivo* situations^14^,^30^. In addition, the miniaturization drastically reduces the Joule heating effect^31^, thereby reducing the chance of any thermotaxis by the cells. Data analysis conducted in this study was similar to Wright et al^38^. The cellular characteristic analyzed in this study includes Forward Migration Index (FMI), where FMI X and FMI Y indicate the efficiency of the forward migration of cells and how they relate to the direction of both axes.

**Fig. 1.**
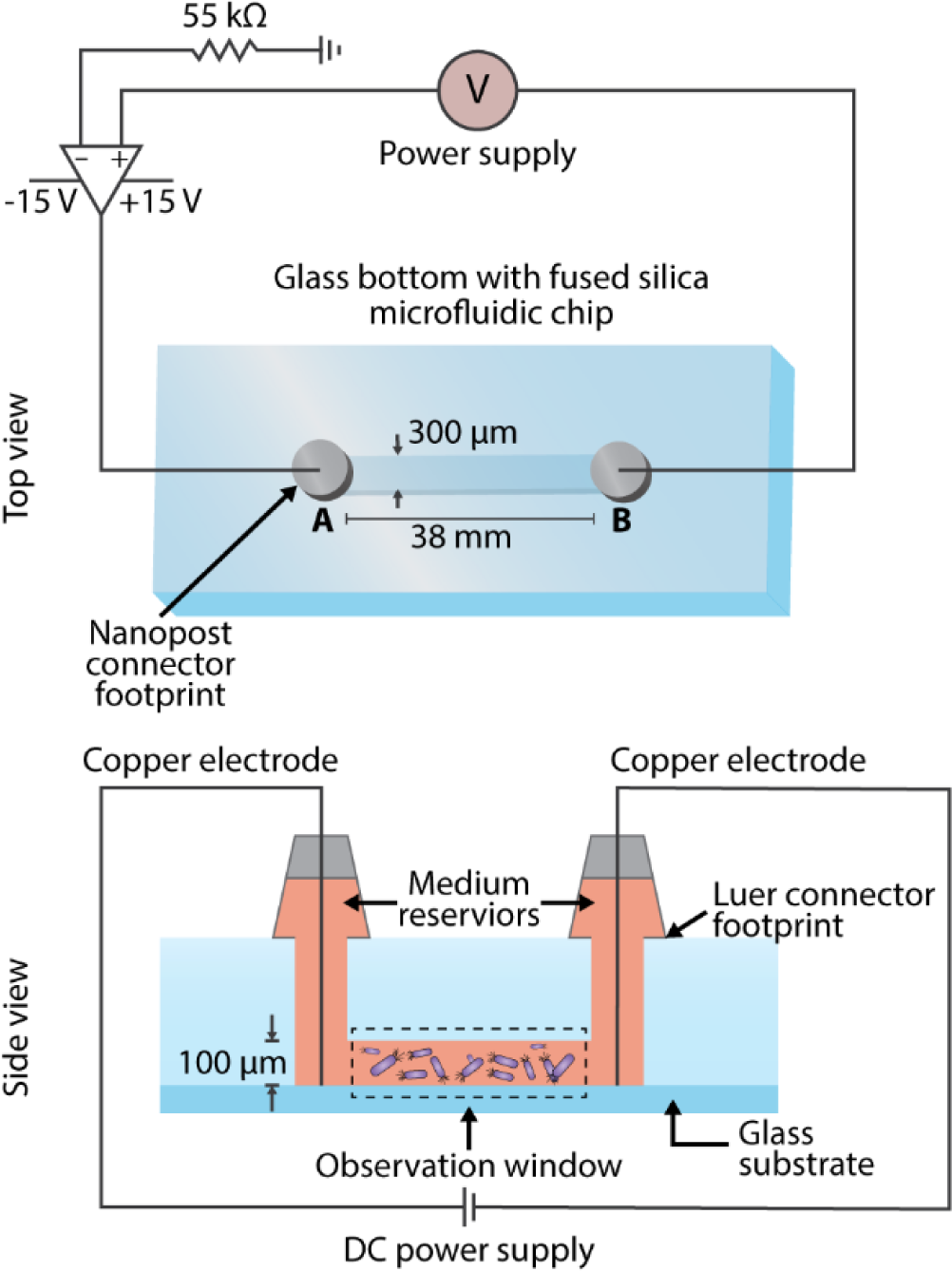
Illustration of the microfluidic experimental setup for bacterial electrotaxis assays. Schematic of the circuit used for generating the desired electric current in the investigation of swimming dynamics of individual bacterial cells.

### Statistical Tracking and Data Analysis

ImageJ (http://rsb.info.nih.gov/ij/) was used to track the cellular frame by frame coordinates, and Ibidi Chemotaxis and Migration Tool software (Ibidi Software, Munich, Germany) was used to calculate the cellular motility metrics of the combined cell tracks. The total number of cell tracks for each setting was 70. At least five independent experiments were carried out. All quantitative data were presented as the mean value±standard deviation. Student’s t-test was applied to compare two distinct groups. P value <0.05 was considered to be statistically significant.

## Results and Disussion

### Single-cue Electrotactic Experimental Metrics

Both *E. coli* and *P. aeruginosa* experience a reduction in cellular speed (Figure 2A, B) and an increase in the forward migration index (FMI) in response to the application of DC, along the chemotactic gradient plotted on the y-axis (Figure 2C). There is also a significant increase in directness with current application (Figure 4A, B), relative to the baseline (0 mA). Resulting average cellular speeds for *E. coli* were 9.7 ± 0.5 μm/s, 5.0 ± 0.4 μm/s, 6.0 ± 0.4 μm/s, and 4.6 ± 0.3 μm/s for 0 mA, 0.07 mA, 0.125 mA, and and 0.175 mA DC, respectively (Figure 3C). Resulting average cellular speeds for *P. aeruginosa* were 44 ± 3 μm/s, 34 ± 2 μm/s, 40 ± 2 μm/s, and 75 ± 3 μm/s for 0 mA, 0.07 mA, 0.125 mA, and 0.175 mA DC, respectively (Figure 3D). Differences in the response to current between *E. coli* and *P. aeruginosa*, as measured by in the change cellular speed, likely reflects differences in their motility and chemo/electrotactic sensing systems^32^. This may help to explain why there is an increase in cellular speed in *P. aeruginosa* at 0.175 mA (relative to baseline), while the application of any current reduces *E. coli* cellular speed. However, 0.07 and 0.125 mA currents reduced the motility of both organisms, indicating that there may be an optimal range of current that will predictably and consistently impact several pathogenic bacterial species.

**Fig. 2.**
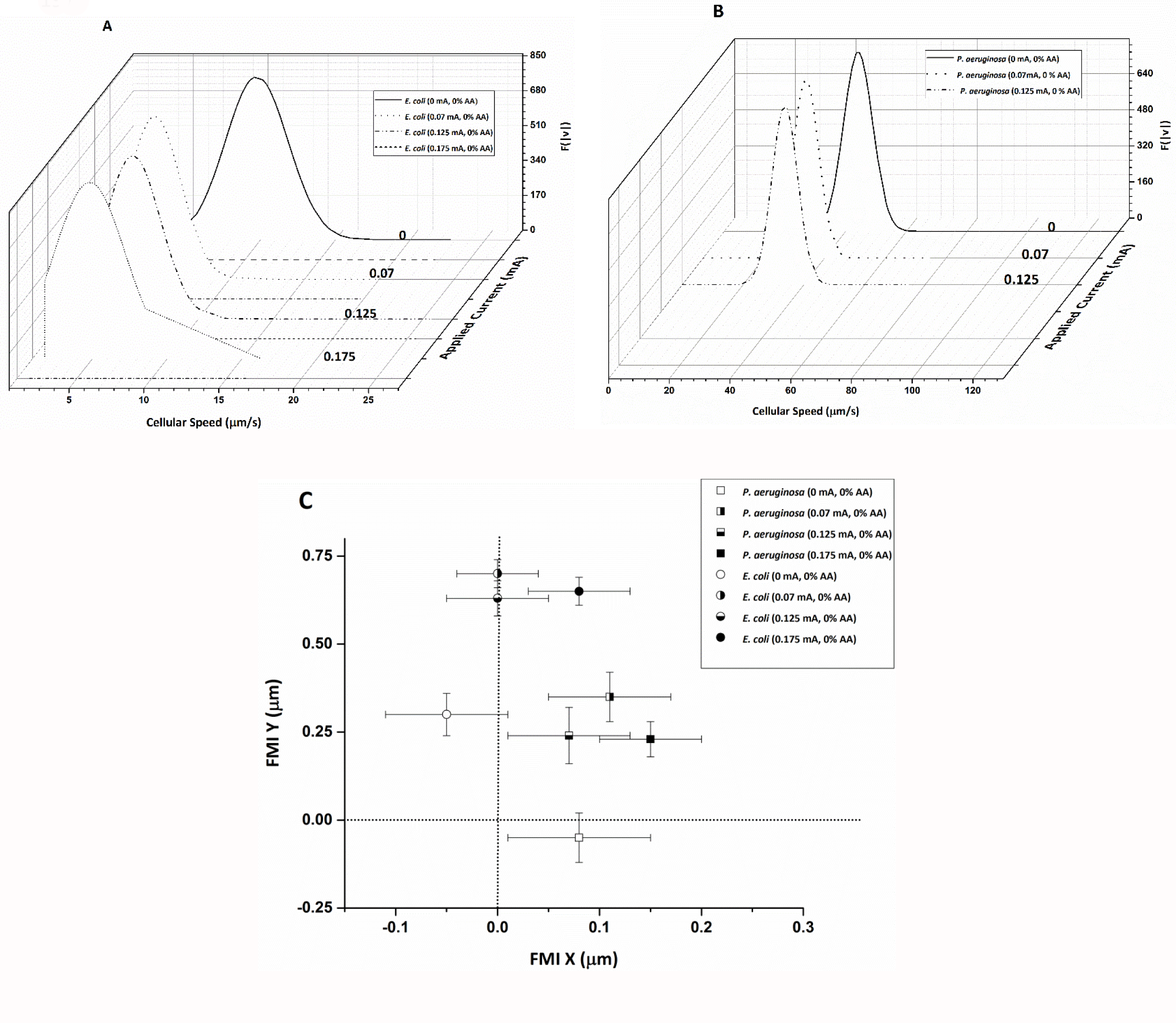
*E. coli* (panel A) and *P. aeruginosa* (panel B) cellular speed distribution in μm/s in response to ES alone at applied current settings listed in mA. The vertical axis indicates resulting scaled Gaussian of distributed cell speeds, F(|**v**|). *E. coli* shows the greatest response to ES alone at 0.07 and 0.175 mA, while *P. aeruginosa* shows the greatest response at 0.07 mA within the standard error. The average forward migration index (FMI) in the x and y directions for *E. coli* and *P. aeruginosa* in response to the application of current is plotted in panel C. Results denote increased migration along the electrotactic gradient (y axis) with the application of current (relative to 0 mA baseline).

**Fig. 3.**
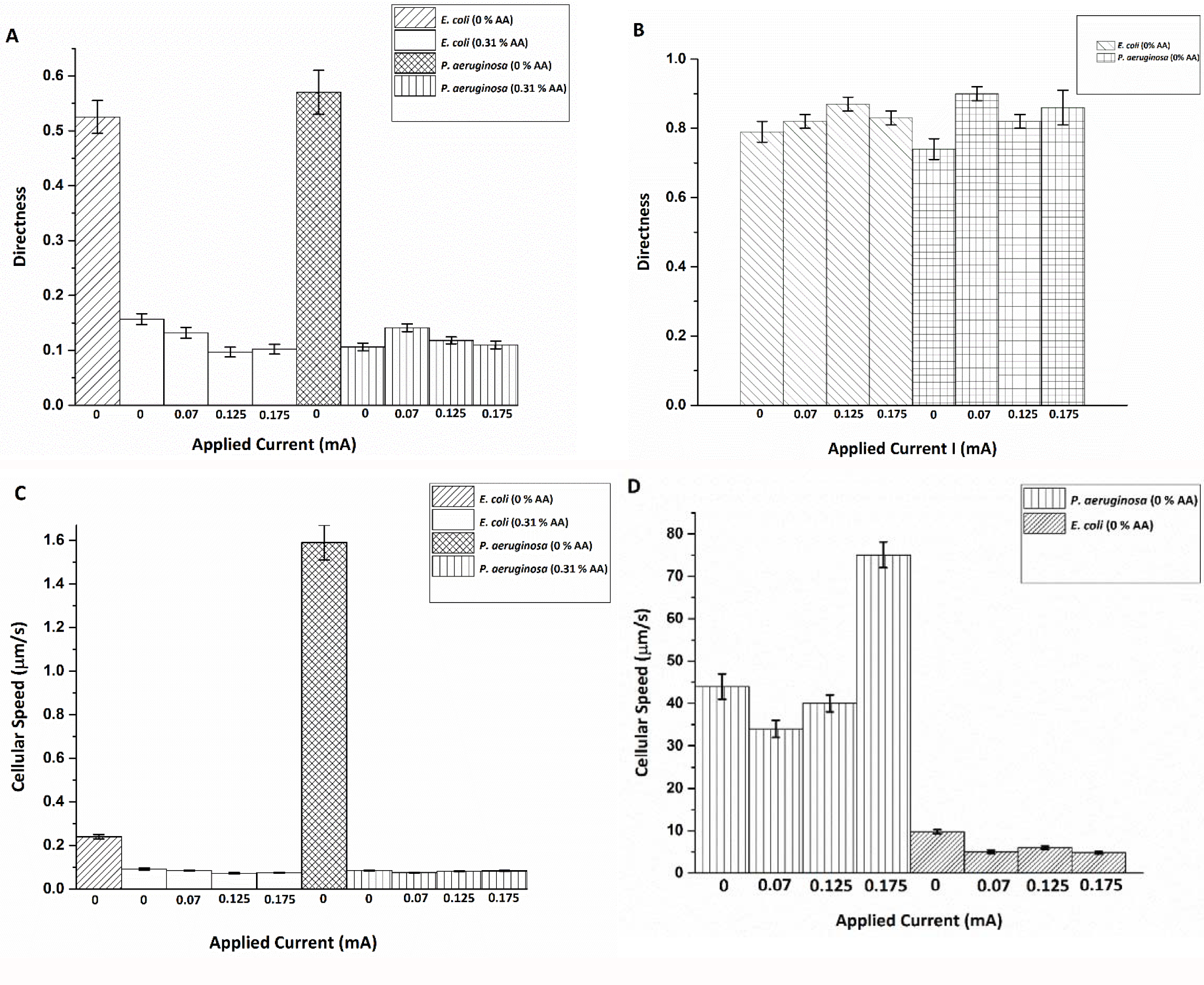
Average directness resulting from applied current and AA concentrations listed for *E*. *coli* (panel A) and *P. aeruginosa* (panel B); average cellular speed for *E. coli* (panel C) and *P. aeruginosa* (panel D). Results indicate an increase in directness relative to baseline with the application of current alone, but a decrease in directness with the application of AA alone or with current. Results also indicate a reduction in cellular speed with 0.31% AA (without or without current) or with current alone, except for *P. aeruginosa* at a current of 0.175 mA (no AA).

### Single-cue Chemotactic Experimental Metrics

Both species experience a reduction in their FMI (Figure 4), average cellular speed (Figure 3 A, B), and directness (Figure 3 C, D) with the introduction of 0.31% AA alone, relative to baseline (no AA). These results indicate that treatment with AA significantly impairs bacterial motility, even when used alone. Furthermore, treatment with AA has equivalent effects on both species, pointing to a broader, more conserved mechanism of action.

**Fig. 4.**
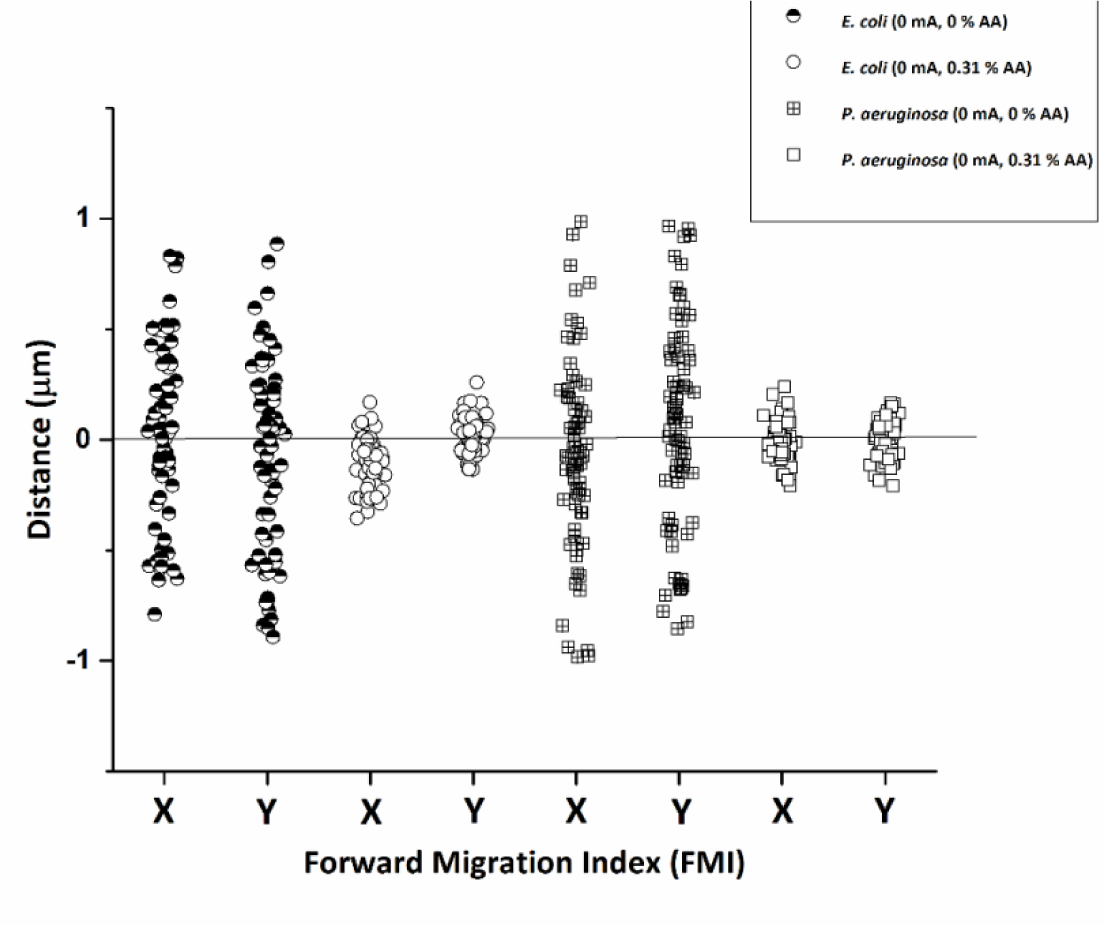
Vertical scatter indicating forward migration index (FMI) distribution in x and y directions for *E. coli* and *P. aeruginosa* comparing the effect of 0.31% AA alone on cells. Results indicate a drastic reduction in migration in both the x and y directions with the application of AA.

Studies have found that bacterial cellular ATP processes are disrupted by exposure to AA^23^, which is one reason why AA has great potential as an antimicrobial agent. Weak acids can cross bacterial membranes more readily than strong acids, because of the equilibrium between their ionized and non-ionized forms, the latter of which can freely diffuse across hydrophobic membranes. This ultimately collapses the proton gradients necessary for ATP synthesis. When acetic acid dissociates, it acidifies the cytoplasm, causing acid-induced proton unfolding and membrane and DNA damage^23^. This effect is specific to bacteria, because host somatic cells contain cholesterol, which controls cell permeability; the interior of the phospholipid bilayer is occupied by hydrophobic fatty acid chains, such that the membrane is impermeable to water-soluble molecules and most biological molecules, including ions^31^.

### Multi-cue Electroceutical Experimental Metrics

Resulting average cellular speeds for *E. coli* in the presence of 0.31% AA were 0.091 ± 0.005, 0.085 ± 0.002, 0.072 ± 0.003, and 0.074 ± 0.002 μm/s for 0 mA, 0.07 mA, 0.125 mA, and 0.175 mA DC, respectively (Figure 3C). Resulting average cellular speeds for *P. aeruginosa* in the presence of 0.31% AA were 0.085 ± 0.002, 0.075 ± 0.002, 0.081 ± 0.001, and 0.083 ± 0.002 μm/s for 0 mA, 0.07 mA, 0.125 mA, and 0.175 mA DC, respectively (Figure 3D).

A reduction in cellular speed relative to baseline (0.31% AA, no current) (Figure 3C, D and Figure 5) was pronounced for both *E. coli* and *P. aeruginosa* upon electroceutical application. The most dramatic reductions in speed occured at 0.070 mA DC for *P. aeruginosa*, while *E. coli* was equally impaired at higher currents (0.125 and 0.175 mA DC). These results suggest that there is an ideal current range in an electroceutical setting for *E. coli*, which would be upwards of 125 μA, while the ideal range for *P. aeruginosa* would be less than 0.125 mA for both single and multi-cue situations (namely 0.07 mA). However, a current of approximately 0.07 mA may be affective across a broader range of strains, meaning that a single application of current (with AA, especially) could provide powerful disinfection in the context of a wound infection. A very pronounced reduction in FMI was also observed at 0.175 mA DC for *E. coli* and at 0.07 mA for *P. aeruginosa* when single-cue ES was compared to our electroceutical approach (AA + current) (Figure 6).

**Fig. 5.**
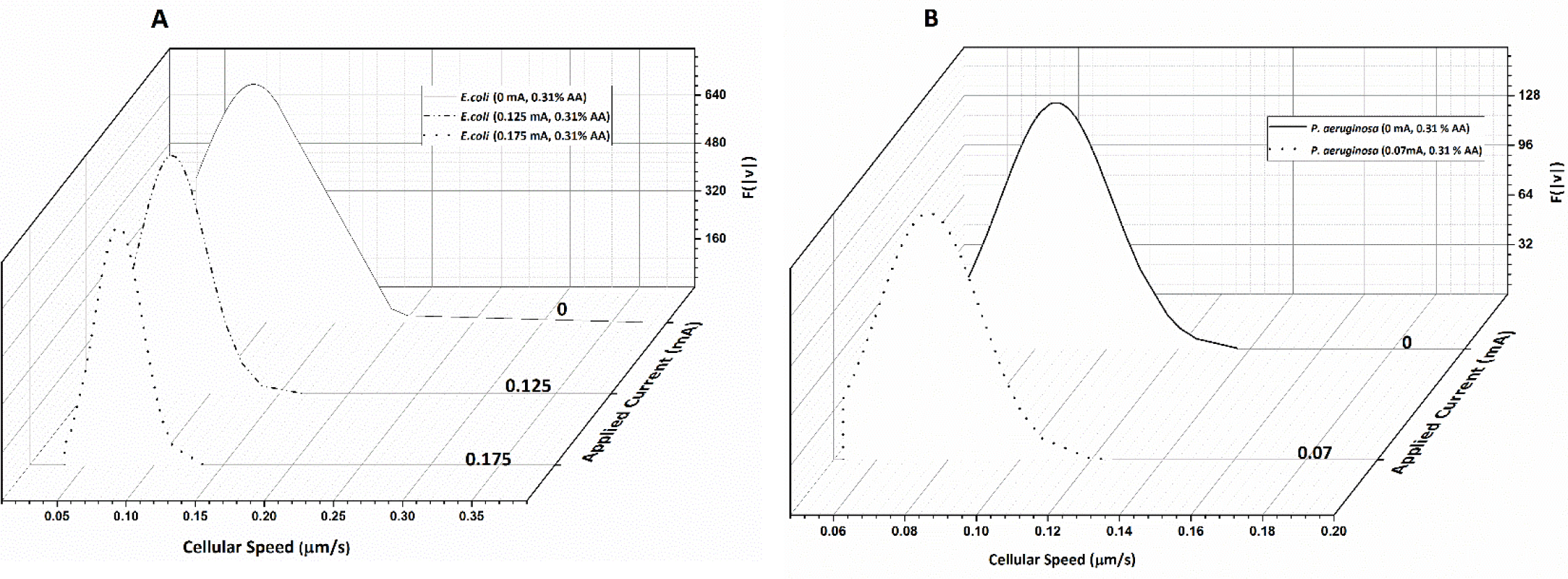
*E. coli* cellular speed distribution in μm/s in response to ES and 0.31% AA specific currents. Results indicate a great reduction in cellular speed upon electroceutical treatment (panel A). *E. coli* was equally responsive to both 0.125 mA and 0.175 mA settings *P. aeruginosa* cellular speed distribution in μm/s (panel B). Results indicate a great reduction in cellular speed with electroceutical treatment of *P. aeruginosa* at the 0.07 mA setting.

**Fig. 6.**
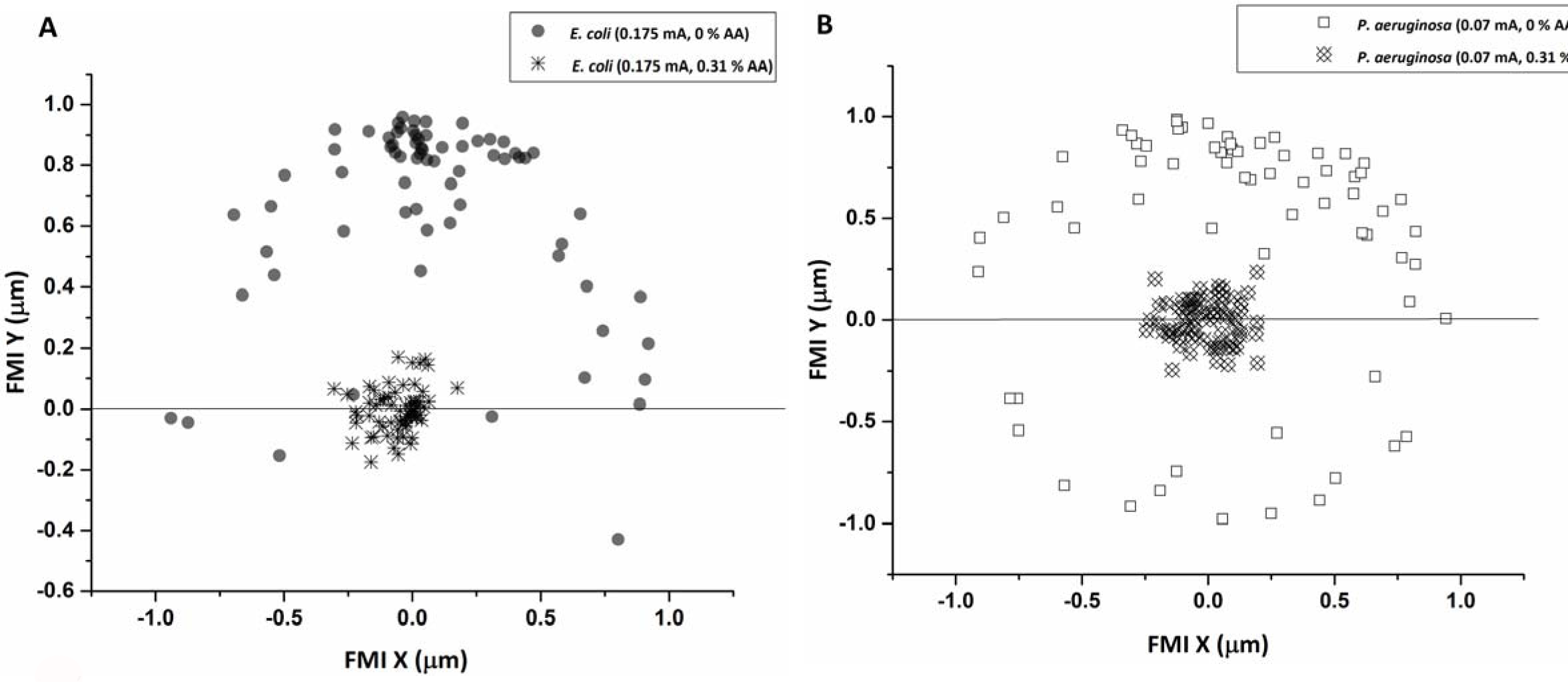
The forward migration index (FMI) distribution for *E. coli* at the 0.175 mA setting in the presence and absence of 0.31% AA (panel A). Results indicate a drastic reduction in overall migration when 0.31% AA was applied in combination with ES at 0.175 mA The FMI distribution for *P. aeruginosa* at a 0.07 mA setting in the presence and absence of 0.31% AA (panel B). Results indicate a drastic reduction in overall migration when 0.31% AA was applied in combination with ES at 0.07 mA.

## Conclusions

Clearly, *P. aeruginosa* responds well to both ES and our electroceutical approach at 0.70 mA DC, while *E. coli* is more responsive at higher current (0.175 mA DC). It is not surprising that *E. coli* and *P. aeruginosa* respond differently to ES settings, as their chemotactic sensing systems differ. *P. aeruginosa* has a more complex sensing system than most microbes with multiple chemotaxis genes that constitute several chemotaxis systems with defined functions^32^. *E. coli* uses a two-component regulatory system consisting of an extracellular sensor and response regulator^32^ that is more susceptible to oxidative stress than *P. aeruginosa*^33^. Hence, electroceutical device design should take into account the differing impacts that such disinfection strategies may have on several relevant wound pathogens. In the case of *P. aeruginosa*, quorum sensing (QS) plays a prominent role in its virulence and biofilm formation, which may offer an ideal target for anti-biofilm therapies.

Oxidative and nitrosative stresses, which can be augmented by electroceutical approaches, play important roles in bacterial inhibition/elimination *in vivo.* In particular, *E. coli* can be toxified by as little as 5 μM extracellular hydrogen peroxide^34^-^35^, and because it is a small uncharged molecule, hydrogen peroxide diffuses across membranes rapidly and has the ability to cause profuse DNA damage when the intracellular concentration rises to 1 μM^27^. *P. aeruginosa*, on the other hand, has developed a multitude of defense mechanisms to tolerate stress conditions, even H_2_O_2_ at relatively high levels, but remains susceptible to other oxidative stressors^33^. Because the buildup of electrochemical oxidative products at the anode and cathode would occur with the application of ES, while ES alone may be a successful strategy for disinfection, but in combination with AA these effects are likely to be augmented. Combining AA with ES should enhance disinfection, as AA has the ability to disrupt essential ATP processes and cause cellular DNA damage, thereby impairing both the ability to communicate and initiate defense mechanisms.

As motility is one microbial defense against opsonization by the host immune defenses, locomotive impairment would effectively trap bacterial cells, allowing for their clearance by migrating phagocytic leukocytes. Since migratory cell behavior in the presence of electrical cues is a naturally existing process within the human body, and if we take the 0.1 μA/mm^2^ DC current density generated as a lower limit for directing leukocyte migration for wound healing, the corresponding current density settings tested here are likely to induce leukocyte migration that would accelerate the natural wound healing process while simultaneously inhibiting opportunistic pathogens. An applied electric field within the physiological range can also induce the directional electrotaxis of epithelial cells and fibroblasts, along with neutrophils and endothelial cells, suggesting a potential role in cellular positioning during wound healing^36^. Because phagocytic neutrophils and monocytes are up to 20 μm in diameter^37^, have cell walls that actively prohibit entry of water soluble molecules^31^, and have a negative surface potential, unlike invading pathogens, they can migrate to the wound site unaffected. As a consequence, our electroceutical approach could create a bacterial trap that would accelerate the natural wound healing process, while augmenting bacterial clearance.

## Acknowledgements

The authors sincerely thank the Natural Sciences and Engineering Research Council of Canada (NSERC), the Ontario Ministry of Agriculture, Food and Rural Affairs (OMAFRA), and the Ontario Ministry of Research and Innovation (MRI) for funding this study.

